# Jasmonate inhibits adventitious root initiation through transcriptional repression of *CKX1* and activation of *RAP2.6L* transcription factor in Arabidopsis

**DOI:** 10.1101/2021.04.23.441150

**Authors:** Asma Dob, Abdellah Lakehal, Ondrej Novak, Catherine Bellini

**Affiliations:** Umeå Plant Science Centre, Department of Plant Physiology, Umeå University, SE-90736 Umeå, Sweden; Laboratory of Growth Regulators, Faculty of Science, Palacký University and Institute of Experimental Botany, Academy of Sciences of the Czech Republic, 78371 Olomouc, Czech Republic; Department of Forest Genetics and Physiology, Umeå Plant Science Center, Swedish Agriculture University, SE-90183 Umea, Sweden; Institut Jean-Pierre Bourgin, INRA, AgroParisTech, CNRS, Université Paris-Saclay, FR-78000 Versailles, France

## Abstract

Adventitious rooting is a *de novo* organogenesis process that enables plants to propagate clonally and cope with environmental stresses. Adventitious root initiation (ARI) is controlled by interconnected transcriptional and hormonal networks, but there is little knowledge of the genetic and molecular programs orchestrating these networks. Thus, we have applied genome-wide transcriptome profiling to elucidate the profound transcriptional reprogramming events preceding ARI. These reprogramming events are associated with downregulation of cytokinin (CK) signaling and response genes, which could be triggers for ARI. Interestingly, we found that CK free-base content declined during ARI, due to downregulation of *de novo* CK biosynthesis and upregulation of CK inactivation pathways. We also found that MYC2-dependent jasmonate (JA) signaling inhibits ARI by downregulating expression of the *CYTOKININ OXIDASE/DEHYDROGENASE1* gene. We also demonstrated that JA and CK synergistically activate expression of *RELATED to APETALA2.6 LIKE* (*RAP2.6L*) transcription factor, and constitutive expression of this transcription factor strongly inhibits ARI. Collectively, our findings reveal that previously unknown genetic interactions between JA and CK play key roles in ARI

## Introduction

Adventitious rooting is a developmental process and adaptive strategy that enables plants to propagate clonally and cope with various environmental stresses (Steffens and Rasmussen, 2016; Lakehal and Bellini, 2019). Like most *de novo* organogenesis processes, adventitious root initiation (ARI) is tightly controlled by coordinated transcriptional networks consisting of complex circuits and feedback loops (Lakehal *et al*., 2019*b*). The underlying genetic and molecular programs are not well understood, but both synergistic and antagonistic hormonal crosstalk is strongly involved in regulation of the transcriptional networks (Bellini *et al*., 2014; Lakehal and Bellini, 2019; Lakehal *et al*., 2020*b*).

Auxin signaling promotes ARI by modulating homeostasis of the negative regulator jasmonate (JA) in Arabidopsis (Sorin *et al*., 2005; Gutierrez *et al*., 2009, 2012). Auxin, perceived by the F-box proteins TRANSPORT INHIBITOR1/AUXIN-SIGNALLING F-BOX PROTEIN (TIR1 and AFB2), triggers degradation of the repressor proteins AUXIN/INDOLE-3-ACETIC ACID 6 (IAA6), IAA9 and IAA17 and subsequently allow AUXIN RESPONSE FACTOR 6 (ARF6) and ARF8 transcription factors to induce expression of GRETCHEN HAGEN3 (GH3.3), GH3.5 and GH3.6 enzymes (Gutierrez *et al*., 2012; Lakehal *et al*., 2019*a*). These three enzymes redundantly promote ARI by conjugating JA to amino acids (Gutierrez *et al*., 2012). Accordingly, genetic analysis has confirmed that MYC2-mediated JA signaling in the xylem-pole pericycle cells represses early ARI events (Lakehal *et al*., 2020*a*). Moreover, JA represses ARI through transcriptional activation of the *APETALA2/ETHYLENE RESPONSE FACTOR115* (*ERF115*) transcription factor and its closely-related paralogs *ERF114* and *ERF113*, also known as *RELATED to APETALA2.6 LIKE/* (*RAP2.6L*) (Lakehal *et al*., 2020*a*). These three transcription factors control a number of developmental programs, including wound healing, shoot and root regeneration, callus formation, stem cell replenishment and ARI (Che *et al*., 2006; Heyman *et al*., 2013, 2016; Ikeuchi *et al*., 2017; Kong *et al*., 2018; Zhou *et al*., 2019; Lakehal *et al*., 2020*a*). The mechanisms enabling these transcription factors to play precise roles in multiple processes are still poorly understood, but they may act through preferential activation or repression of downstream targets in a context-dependent manner. For example, *ERF115* inhibits ARI by promoting *de novo* cytokinin (CK) biosynthesis through transcriptional regulation of the *ATP/ADP ISOPENTENYLTRANSFERASES 3* (*IPT3*) gene (Lakehal *et al*., 2020*a*). *IPT3* encodes an enzyme that catalyzes the first and rate-limiting step in *de novo* CK biosynthesis (Miyawaki *et al*., 2006). CKs are adenine-derived phytohormones that control a plethora of developmental processes including adventitious rooting (Kieber and Schaller, 2014). Like JA, CKs inhibit ARI in a dose-dependent manner and their action seems to be evolutionary conserved among various plant species (Ramírez-Carvajal *et al*., 2009; Mao *et al*., 2019; Lakehal *et al*., 2020*a*). Thus, previous studies have provided substantial information about roles of JA and CKs, but not the interplay between them during ARI, or even if they directly interact during the process. To acquire such information, we have examined their interactions. As reported here, we found that dark to light shifts decrease CK free-base contents and hence repress CK signaling outputs. We also show that JA inhibits ARI through transcriptional repression of the *CKX1* gene. Moreover, JA and CK additively induce expression of the *RAP2.6L* transcription factor, which is a negative regulator of ARI.

## Results

### Dark-light transition triggers profound transcriptional reprogramming

To uncover the transcriptional reprogramming events associated with ARI in Arabidopsis we re-analyzed our publicly-available transcriptome dataset (Lakehal *et al*., 2020*a*). This dataset was generated by sequencing RNA extracted from etiolated hypocotyls of wild-type seedlings (Colombia-0 ecotype, Col-0) at T0 (just before moving them into light), at T9 (9 h after the transfer) and T24 (24 h after the transfer) (Fig. 1A). This analysis detected 3778 differentially expressed genes (DEGs), 2480 of which were upregulated and 1298 downregulated at T9 compared to T0 (Fig. 1B and Supplementary Table S1). It also detected 4709 DEGs between T0 and T24, 3001 of which were upregulated and 1708 downregulated at T24 compared to T0. Moreover, 859 were differentially expressed (433 upregulated and 426 downregulated) at T24 compared to T9 (Fig. 1B and Supplementary Table S2-S3). These data clearly show that shifting such seedlings from dark to light causes profound transcriptional reprogramming in the hypocotyls, which likely involves changes in expression of genes that regulate cell-fate decision programs leading to ARI.

**Figure 1.**
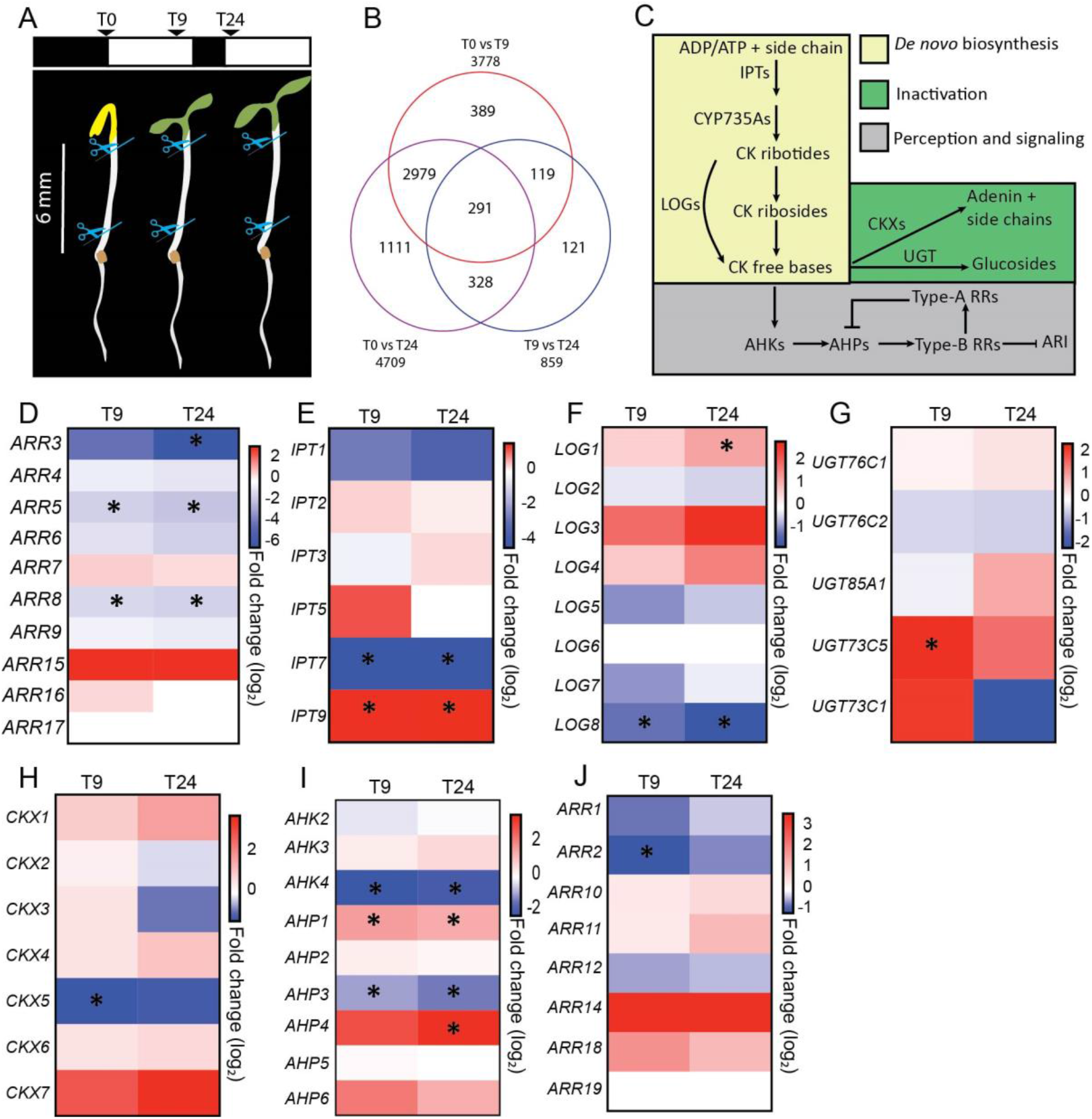
Dark-light transition affects CK pathways. **(A)** Representative scheme of the experimental setup used to collect samples for RNA sequencing. **(B)** Venn diagram showing numbers of differentially expressed genes during ARI in wild type (Col-0) seedlings. **(C)** Illustrative model **of** CK pathways. **(D-J)** Heatmaps of the genes involved in the CK pathways: **(D)** response, **(E-F)** *de novo* biosynthesis, **(G)** inactivation, **(H)** irreversible cleavage and **(I-J)** signaling. Heatmaps represent fold-changes (log2) in transcript abundance in wild type seedlings. Blue and red colors indicate downregulated and upregulated expression, respectively, at T9 or T24 (9 and 24 h after transfer to light, at T0 respectively), relative to T0. Asterisks indicate significant differences.

### CK pathways are reprogrammed during early stages of ARI

We have previously shown that CKs repress ARI in Arabidopsis (Lakehal *et al*., 2020*a*), but the mechanisms modulating CK pathways during this process have remained unclear. To clarify them, we first checked expression profiles of type-A *ARABIDOPSIS RESPONSE REGULATOR* (*ARR*) genes, which are used as proxies for CK response status (Fig. 1C) (Brenner *et al*., 2005; Bhargava *et al*., 2013; Kieber and Schaller, 2018). Interestingly, we found that three type-A *ARR* genes (*ARR3, ARR5* and *ARR8*) were downregulated upon shifting seedlings from dark to light, suggesting that CK signaling is repressed during early stages of ARI (Fig. 1C-1D). To understand causes of the repression of CK signaling, we examined expression profiles of genes involved in CK homeostasis and signaling pathways (Kieber and Schaller, 2014, 2018) (Fig. 1C). We found that several key genes were downregulated upon shifting seedlings from dark to light. These included *IPT7* and *LONELY GUY* (*LOG8*) involved in *de novo* CK synthesis, *ARABIDOPSIS HISTIDINE KINASE 4 (AHK4*) and *HISTIDINE-CONTAINING PHOSPHOTRANSMITTER3* (*AHP3*) involved in perception and transduction, and type-B *ARR2* involved in transcriptional regulation. In addition, *UDP-GLUCOSYL TRANSFERASE 73C5* (*UGT73C5*), encoding an enzyme catalyzing CK inactivation (Hou *et al*., 2004), was upregulated (Fig. 1E-1J and Supplementary Tables S1-S3). These results suggest that repression of CK signaling may be due to reduction in CK biosynthesis and increase in CK inactivation. However, we also found that *IPT9, LOG1, AHP1* and *AHP4* were upregulated and *CYTOKININ OXIDASE/DEHYDROGENASE5* (*CKX5*) downregulated upon shifting the seedlings from dark to light, suggesting the possible existence of compensatory mechanisms (Fig. 1E-1I and Supplementary Tables S1-S3).

Taken together, these results suggest that dark to light transition generates cues that repress CK signaling and thereby de-repress the gene expression programs leading to ARI.

### Dark-light transition decreases *de novo* biosynthesis of CKs and increases their inactivation

As dark-light transition led to a repression of CK responses that was coupled with downregulation of genes involved in *de novo* biosynthesis and upregulation of a gene involved in CK inactivation, we hypothesized that downregulation of CK signaling is caused by reduction in CK free-base content. To test this hypothesis, we quantified the CK free-bases: isopentenyladenine (iP), *trans-*Zeatin (*t*Z), *cis-*Zeatin (*c*Z) and dihydrozeatin (DHZ). We also quantified their riboside precursors (iPR, *t*ZR, *c*ZR, DHZR), corresponding 5-monophosphate ribotides (iPRMP, *t*ZRMP, *c*ZRMP, DHZMP) and glucosyl conjugates: CK *N*-glucosides (iP7G, *t*Z7G, *c*Z7G, DHZ7G, iP9G, *t*Z9G, *c*Z9G, DHZ9G) and CK *O*-glucosides (*t*ZROG, *t*ZOG, *c*ZROG, *c*ZOG, DHZROG, DHZOG). Glucosyl conjugates (*N*-glucosides and *O*-glucosides) are inactive and the CK *N*-glucosides are even thought to be irreversibly inactive (Kieber and Schaller, 2014), except for tZ7G and tZ9G, which may be reactivated by conversion to *t*Z (Hošek *et al*., 2020). The CK contents of the seedlings were quantified under the same conditions and at the same time points as RNA in the RNA sequencing experiments (Fig. 1A), except T72 (72 h after transferring etiolated seedlings to light) instead of T24 to cover a wider developmental window of AR development. We found that tZRMP, tZR, tZ, iPR and iP contents all declined after moving the seedlings to light (Fig. 2A-2C). We also observed slight decreases in cZRMP and cZ at T72 compared to T0 (Fig. 2A-2C). Moreover, the dark-light transition increased the conjugation process, as manifested by increases in iP7G, iP9G tZ7G, tZ9G, cZ9G, tZOG, tZROG, cZOG, and cZROG (Fig. 2D-2F). Interestingly, we also observed reductions in DHZR and DHZ, accompanied by increases in DHZ7G, DHZ9G and DHZOG after moving the seedlings to light (Supplementary Fig. 1A-1E). Thus, although their biological significance and exact role are not yet well understood, it seems that DHZ-type CKs might also play a role in ARI.

**Figure 2.**
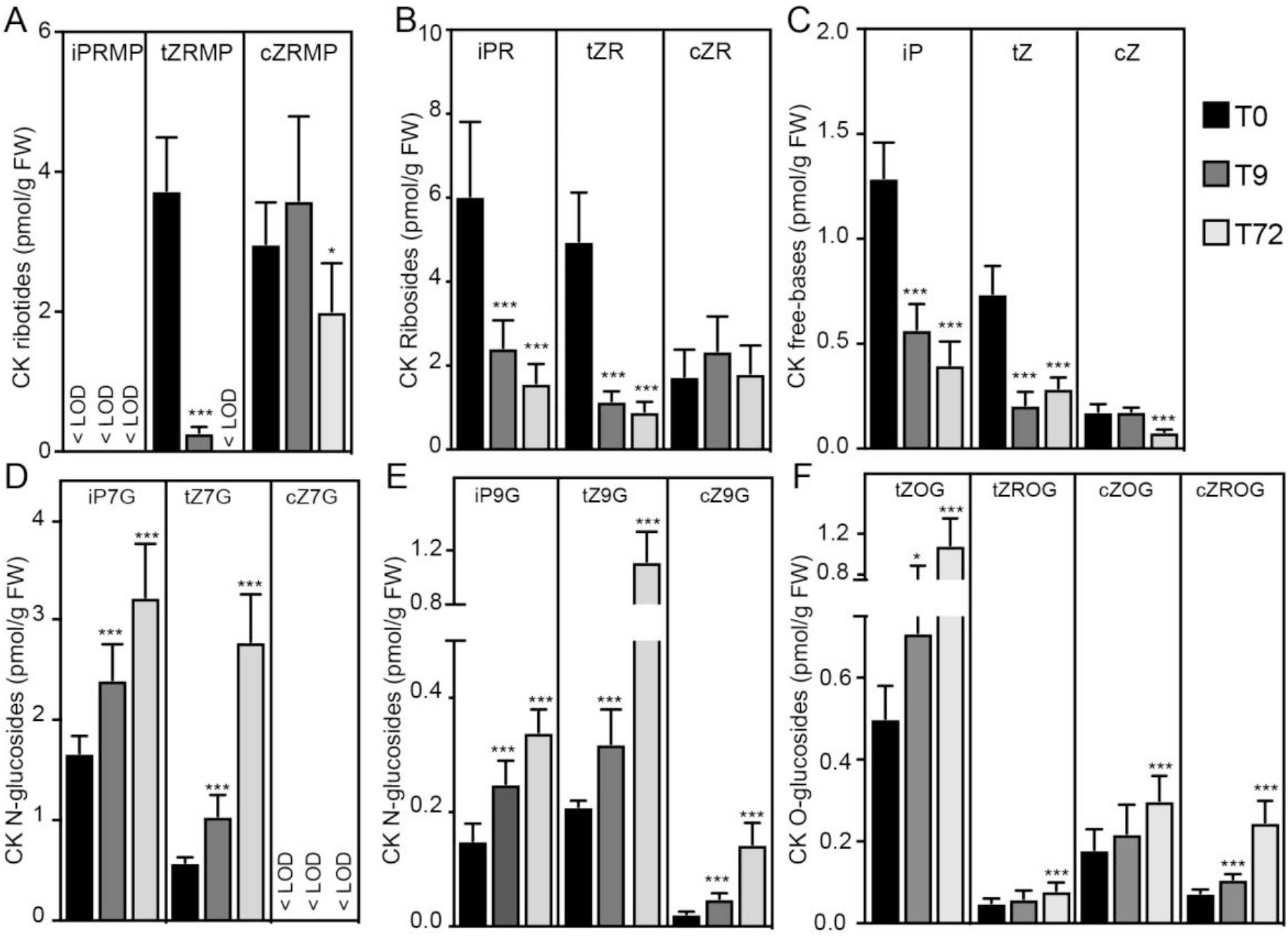
Dark-light transition affects CK homeostasis. Contents in wild type (Col-0) seedlings grown in the dark until their hypocotyls were 6-7 mm long (T0) then shifted to the light for either 9 h (T9) or 72 h (T72) of: **(A)** CK ribotides, **(B)** CK ribosides, **(C)** CK freebases, **(D-E)** CK *N*-glucosides **(F)** CK *O*-glucosides. *, **, and *** indicate significant differences (0.05 > *p* > 0.01, 0.01 > *p* > 0.001, and *p* < 0.001, respectively) at T9 and T72 relative to T0 according to Analysis of Variance with t-tests. Means (bars) and standard deviations (whiskers). <LOD means under the limit of detection.

Collectively, these data indicate that dark-light transition has a dual effect on CK homeostasis, both repressing the *de novo* CK biosynthesis pathways and enhancing the inactivation pathways. Repressing CK-free bases below a certain threshold likely triggers the ARI process and subsequently allows AR primordia development.

### JA downregulates *CKX1* expression in a COI1-dependent manner

Previous studies have shown that moving seedlings from dark to light reduces JA contents of etiolated hypocotyls (Gutierrez *et al*., 2012; Lakehal *et al*., 2020*a*). They have also shown that both JA and CKs repress ARI (Ramirez-Carvajal *et al*., 2009; Gutierrez *et al*., 2012; Mao *et al*., 2019; Lakehal *et al*., 2020*a*), but not whether they directly interact during this process. In this study we found that light to dark transition reduces CK contents of etiolated hypocotyls (Fig. 2A-2F and Supplementary Fig. 1A-1E) as well as their JA contents, raising the possibility that interplay between their pathways provides a coherent developmental input for ARI. To test the hypothesis that JA might control CK content, we first searched publicly-available gene expression datasets (using Arabidopsis eFP browser) for CK-related genes that are transcriptionally affected by exogenous applications of JA or the JA derivative methyl-jasmonate (MeJA) (Winter *et al*., 2007). A particularly interesting finding is that *CKX1* expression was weaker in Arabidopsis seedlings treated with MeJA for 1 h or 3 h than in mock-treated counterparts (Supplementary Fig. 2). *CKX1* encodes an endoplasmic reticulum-localized enzyme that catalyzes irreversible degradation of CKs by cleavage of their side chains (Werner *et al*., 2003; Niemann *et al*., 2018). Interestingly, the *CKX1* gene was slightly upregulated in hypocotyls of seedlings shifted from dark to light, in accordance with their reduction in content of CK free-bases (Fig. 1H). This expression profile was further confirmed by quantitative real-time PCR (qRT-PCR) (Supplementary Fig. S3). These data prompted us to hypothesize that MYC2-dependent JA signaling might control the irreversible cleavage of CKs and thus repress ARI. To test this hypothesis, we first quantified relative amounts of *CKX1* transcripts in wild type seedlings treated with 2, 10 or 20 μM of JA for 1 h. Our data revealed that exogenous applications of JA downregulated expression levels of *CKX1* in a dose-dependent manner (Fig. 3A) and this downregulation requires presence of a functional JA receptor, CORONATINE INSENSITIVE1 (COI1) (Fig. 3B), which triggers transcriptional changes regulated by MYC2.

**Figure 3.**
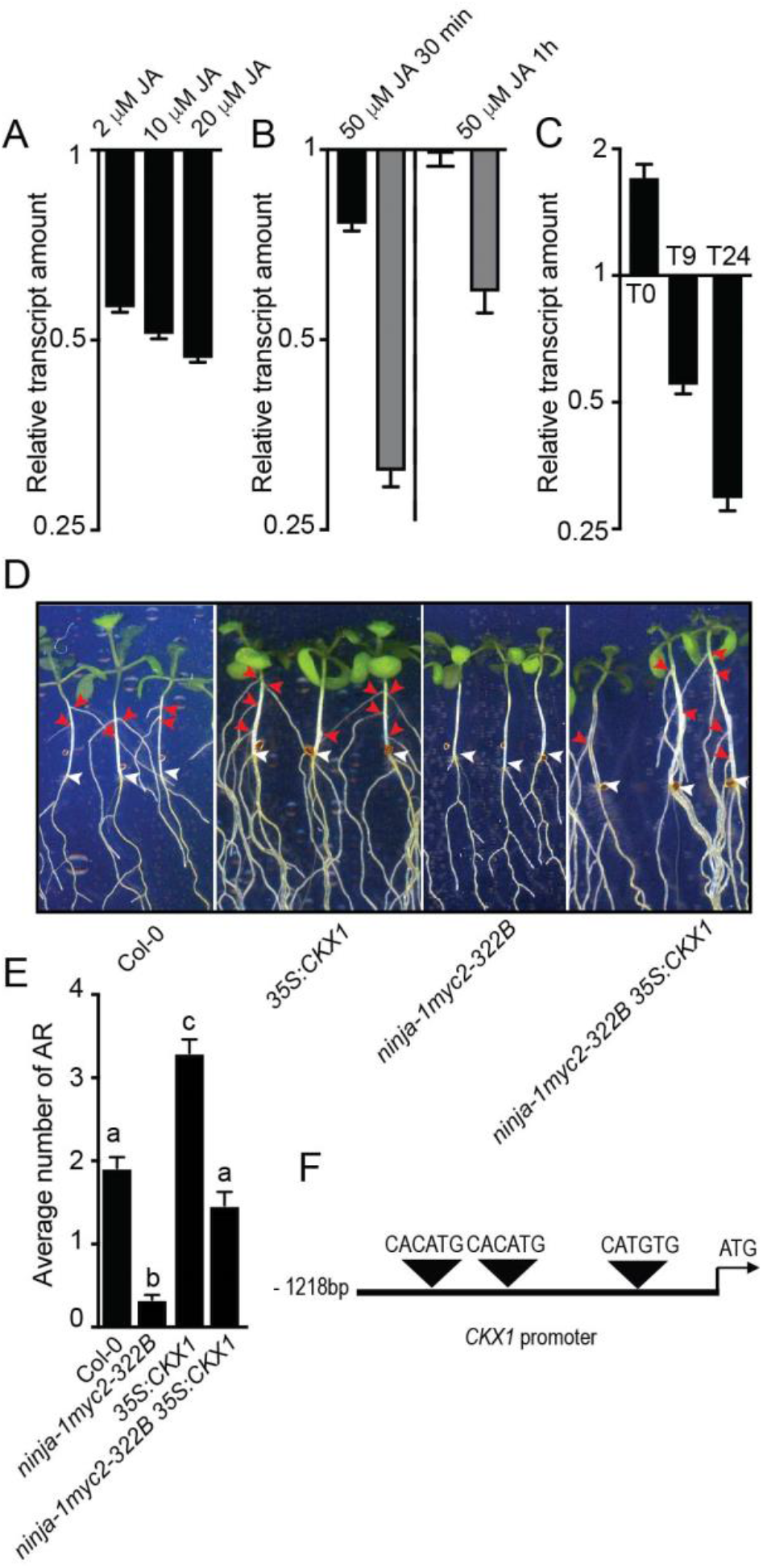
Jasmonate (JA) inhibits adventitious root initiation by repressing *CKX1* expression. **(A-C)** Relative amounts of *CKX1* transcripts measured by qRT-PCR. Means and standard errors of the mean (indicated by error bars) obtained from three technical replicates. All the qRT-PCR experiments were repeated with at least one other independent biological replicate and gave similar results. **(A)** Transcripts extracted from 6-day-old wild-type (Col-0) seedlings treated with JA (at indicated concentrations) for 1 h relative to amounts in mock-treated controls. **(B)** Transcripts extracted from 6-day-old *coi1-16* mutant (dark bars) or wild-type (gray bars) seedlings treated with 50 μM JA relative to amounts in mock-treated controls. **(C)** Transcripts extracted from dissected hypocotyls of *ninja-1myc2-322B* double mutant or wild-type seedlings etiolated in the dark until their hypocotyls were 6 mm long (T0) then moved to light for 9 h (T9) or 24 h (T24) relative to amounts in wild-type (Col-0) seedlings. **(D)** Representative images showing emerged ARs. Red and white arrowheads indicate emerged ARs and root-hypocotyl junctions, respectively. **(E)** Average numbers of ARs. The non-parametric Kruskal-Wallis test combined with Dunn’s multiple comparison post-test indicates that the *ninja-1myc2-322B35S:CKX1* triple mutant produced significantly more AR than the *ninja-1myc2-322B* double mutant. Bars and whiskers indicate means and standard errors of the means, respectively (*n*>50; *P* < 0.0007). **(F)** Illustrative scheme representing locations of G-box-like motifs in the *CKX1* promoter.

To obtain more evidence of MYC2’s involvement, we compared amounts of *CKX1* transcripts in *ninja-1myc2-322B* double mutant and wild type seedlings. *Ninja-1* is a loss-of-function mutation (Acosta *et al*., 2013), whereas *myc2-322B* is a gain-of-function mutation (Gasperini *et al*., 2015). Combination of these two mutations results in constitutive and enhanced JA signaling (Gasperini *et al*., 2015; Lakehal *et al*., 2020*a*). We found that *CKX1* expression was weaker in *ninja-1myc2-322B* double mutants than in wild type seedlings at T9 and T24, but not T0 (Fig. 3C). As JA signaling is constitutively enhanced in *ninja-1myc2-322B* double mutants we expected to observe *CKX1* downregulation at all sampling time points, but this was not the case at T0. These data suggest that the transcriptional regulation of *CKX1* by MYC2-dependent JA signaling involves other light-dependent factor(s), which require further elucidation.’

### *CKX1* action downstream of *MYC2-*dependent JA signaling promotes ARI

To test whether *CKX1* activity downstream of MYC2-dependent JA signaling promotes ARI genetically, we generated triple *35S:CKX1ninja-1myc2-332B* mutants. Double *ninja-1myc2-322B* mutants produced fewer, and overexpressing *35S:CKX1* mutants more, ARs than wild type seedlings (Fig. 3D-3E). These results are consistent with our previous findings (Lakehal *et al*., 2020*a*). Interestingly, the triple *35S:CKX1 ninja-1myc2-332B* mutants produced similar numbers of ARs to the wild type seedlings (Fig. 3D-3E), indicating that enhancing the irreversible cleavage of CKs in the *ninja1myc2-332B* double mutants is sufficient to suppress the negative effect of JA signaling on ARI. The MYC2 transcription factor preferably binds to G-Box or G-box-like *cis*-regulatory elements in the induction or repression of its downstream targets (Godoy *et al*., 2011). Therefore, we searched for such *cis*-regulatory motifs in the 1.5 Kb sequence upstream of the CKX1 gene’s *ATG* translation start codon. Interestingly, we found three G-box-like tetrameric motifs (CACATG, CACATG, CATGTG), suggesting that MYC2 might repress *CKX1* expression by directly binding to its promoter during ARI (Fig. 3F).

### ARI repression by JA and CKs involves synergistic induction of *RAP2.6L*

To obtain insights into the mechanism whereby JA-CK interplay represses ARI, we searched our transcriptomic database for potential transcription factors that are differentially expressed during early stages of the process (Fig. 1A). This analysis detected 180 differentially expressed transcription factors, 91 of which were upregulated and 79 downregulated at T9 compared to T0 (Supplementary Table S4). It also detected 230 differentially expressed transcription factors between T0 and T24, 143 of which were upregulated and 87 downregulated at T24 compared to T0. Moreover, 63 were differentially expressed (39 upregulated and 24 downregulated) at T24 compared to T9 (Supplementary Table S5-S6).

Interestingly, we retrieved several transcription factors with potential role in *de novo* organogenesis and cell stem cell regulations, one of which (*RAP2.6L*) was downregulated at T9 and T24 (Supplementary Tables S4-S6). How *RAP2.6L* is transcriptionally regulated during ARI has remained unclear. We therefore tested whether its expression is affected by exogenous applications of JA and/or CKs. We quantified the relative abundance of this gene’s transcripts in wild type seedlings treated with 25 μM JA, 1uM or 10 μM *t*Z, 25 μM JA + 1uM *t*Z, or 25 μM JA + 10 μM *t*Z. We found that exogenous applications of both 25 μM JA and 10 μM *t*Z slightly induced expression of *RAP2.6L* (Fig. 4A). Moreover, combination of JA 25 μM with either 1 or 10 μM *t*Z had an additive effect on *RAP2.6L* levels (Fig. 4A). These findings suggest that JA and CK synergistically induce expression of *RAP2.6L*. To further investigate whether enhancing *RAP2.6L* expression is sufficient to repress ARI, we analyzed the AR phenotype in transgenic lines overexpressing *RAP2.6L* (*35S:RAP2.6L*). We found that *35S:RAP2.6L* plants produced dramatically fewer AR than wild type (Wassilewskija-4, Ws-4) counterparts (Fig. 4B and 4F). *RAP2.6L* seems to specifically control ARI because LR number and LR density were not affected, although primary roots of *35S:RAP2.6L* seedlings were slightly longer than those of wild type counterparts (Fig. 4C-4E).

**Figure 4:**
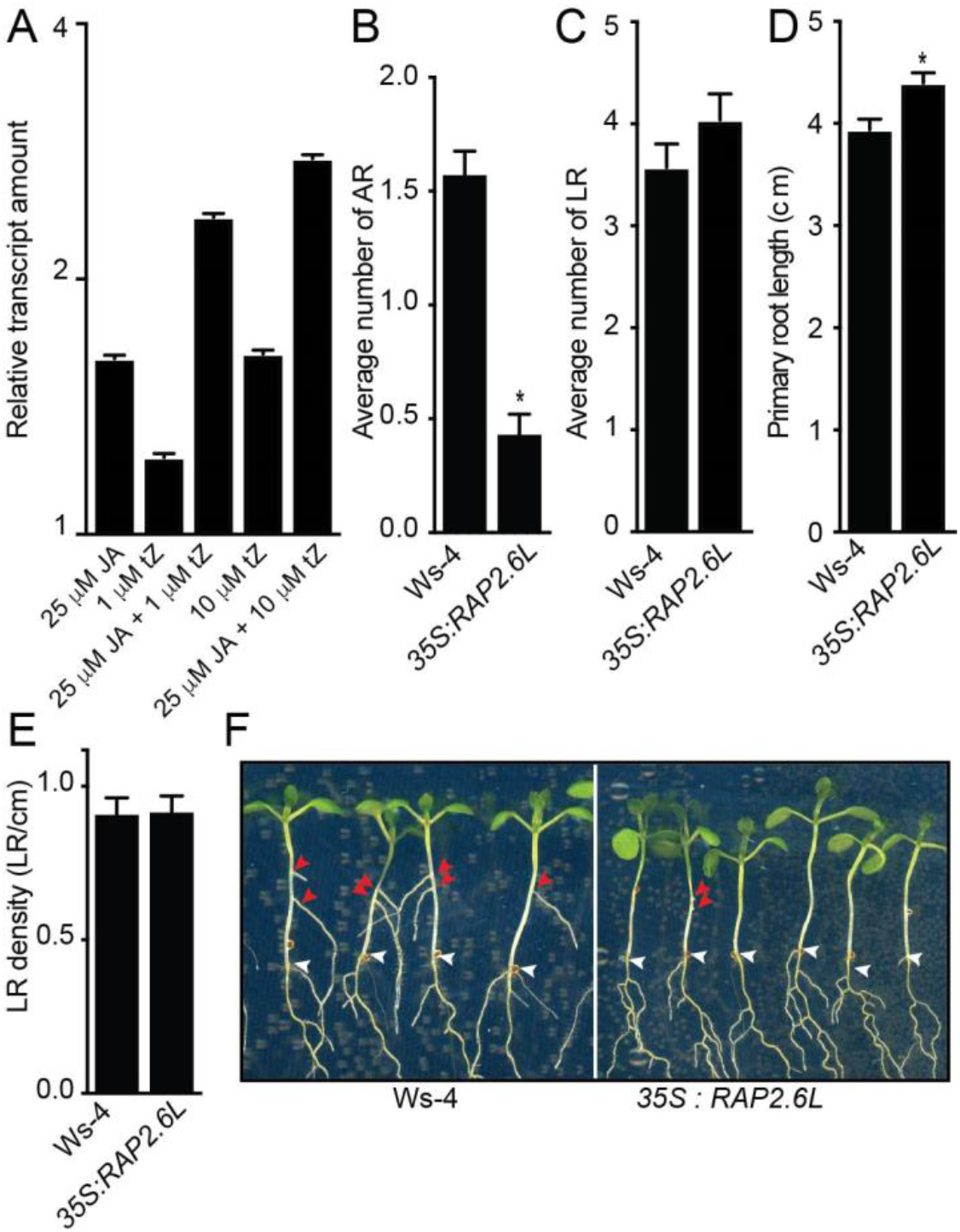
CK and JA synergistically repress ARI through *RAP2.6L* induction. **(A)** Amounts of *RAP2.6L* transcripts in wild type seedlings treated with indicated concentrations of JA and/or *t*Z relative to amounts in mock-treated controls. Means and standard errors of the mean obtained from three technical replicates. The qRT-PCR experiment was repeated with another independent biological replicate and gave similar results. **(B)** Average numbers of ARs. The t-test indicates that *35S:RAP2.6L* produced significantly fewer AR than wild type seedlings (*n*>60; *P* < 0.0001). **(C)** Average numbers of LRs. ‘(D and E) Results of measurements respectively showing that *35S:RAP2.6L* mutants produced longer primary roots than wild type seedlings (t-test: *n*>24, *P* < 0.0022) but their lateral root density did not differ **(B-E)** Bars and whiskers indicate means and standard errors of the means, respectively. **(F)** Representative images showing numbers of emerged ARs. Red and white arrowheads indicate emerged ARs and junction roots, respectively.

We concluded that CKs and JA may repress ARI by synergistically inducing expression of *RAP2.6L*.

## Discussion

CKs control a plethora of developmental processes including ARI (Wybouw and De Rybel, 2019). They also mediate plant responses to a wide spectrum of environmental cues (Kieber and Schaller, 2014). These diverse functions involve tight spatiotemporal control of dynamic CK pools, especially the CK free bases, which are mainly governed by the combined action of three interconnected processes: biosynthesis, conjugation and cleavage (Sakakibara, 2006; Kieber and Schaller, 2014, 2018). Moreover, CKs’ precise roles in physiological and developmental processes involve complex crosstalk with other hormones, which adds further regulatory layers (O’Brien and Benková, 2013). Previous studies have firmly established that CKs play a key role in ARI, but little knowledge of the mechanisms modulating their pools during early stages of the process.

In the study reported here we found that dark-light shifts cause profound transcriptional reprogramming of CK pathways in etiolated seedlings, resulting in substantial reductions in CK contents and signaling that are likely triggers of ARI events. Accordingly, reducing CK contents in Arabidopsis either through knocking out key enzymes for their *de novo* biosynthesis (e.g. *ipt3ipt5ipt7*) or enhancing their irreversible cleavage (as in the *35S:CKX1* overexpressing line) reportedly leads to significantly more ARs than in wild type seedlings (Werner *et al*., 2003; Miyawaki *et al*., 2006; Lakehal *et al*., 2020*a*). Blocking the CK perception or signaling pathways (as in *ahk3ahk4, arr1arr11* and *arr1arr11arr12* mutants) leads to similar increases in AR number (Rasmussen *et al*., 2012; Lakehal *et al*., 2020*a*).

Interactions between light and phytohormones have already been proposed, but no clear mechanistic link between light and CKs has been established, and further research is needed to identify the light-related genes mediating the interactions (if any) involved. However, light modulates the homeostasis and signaling of several hormones including auxin (Halliday *et al*., 2009), which has antagonistic regulatory effects to CKs in diverse developmental programs including ARI (Schaller *et al*., 2015; Lakehal and Bellini, 2019). As auxin may putatively repress CK biosynthesis (Nordström *et al*., 2004; Cheng *et al*., 2013; Zhang *et al*., 2018), we are tempted to speculate that the increase in auxin content and upregulation of ARF6- and ARF8-dependent auxin signaling observed after shifting etiolated seedlings from dark to light (Tian *et al*., 2004; Gutierrez *et al*., 2012) could directly impair CK biosynthesis. Another plausible hypothesis is that auxin signaling controls CK content through JA (Gutierrez *et al*., 2012; Lakehal *et al*., 2020*a*). Accordingly, we found that downregulation of CK content coincided with a reduction in JA content, supporting a link between the two hormone pathways. Our gene expression data suggested that MYC2-mediated JA signaling represses ARI through transcriptional repression of *CKX1*, which catalyzes irreversible cleavage of CKs (Werner *et al*., 2003). The transcriptional repression of *CKX1* would inevitably lead to accumulation of CKs, in the absence of reductions in their synthesis. This is consistent with previous finding that exogenous applications of MeJA led to a transient and rapid accumulation of CKs in *Triticum aestivum* L. (wheat) seedlings (Avalbaev *et al*., 2016), which was attributed to MeJA having a dual role, both downregulating *CKX1* expression and impairing enzymatic activity of its product (Avalbaev *et al*., 2016).

There have been few demonstrations of interplay between JA and CKs. However, it has been recently shown that CK signaling promotes JA accumulation through transcriptional activation of key genes in the JA biosynthesis pathway and thus inhibits leaf growth in *Zea mays* (Uyehara *et al*., 2019). In the context of ARI, the possibility that CK might promote JA biosynthesis and/or signaling via a positive feedback loop cannot be excluded. Interestingly, our data revealed that JA and CK additively promote expression of the *RAP2.6L* transcription factor, which is a negative regulator of ARI. These findings indicate that the crosstalk between JA and CK is complex and might involve parallel pathways that require further elucidation. *RAP2.6L* gene belongs to *APETALA2/ETHYLENE RESPONSE FACTOR* subfamily X (Nakano *et al*., 2006; Heyman *et al*., 2018) and has been implicated in a number of developmental and regenerative processes, including shoot meristem proliferation and regeneration as well as tissue-reunion (Che *et al*., 2006; Yang *et al*., 2018). (Che *et al*., 2006) found that the *RAP2.6L* gene is strongly upregulated in root explants incubated in cytokinin-rich shoot-inducing medium and shoot regeneration is severely impaired in *rap2-6l* mutants under these conditions, indicating that *RAP2-6L* is required for CK-mediated shoot regeneration. *RAP2.6L* is also reportedly upregulated at sites of wounds on inflorescence stems, together with *LIPOXYGENASE2*, which encodes an enzyme involved in JA biosynthesis, suggesting that expression of *RAP2.6L* may be triggered by wound-induced JA (Asahina *et al*., 2011). The cited authors also found that exogenous applications of MeJA upregulated *RAP2.6L* expression (Asahina *et al*., 2011), in accordance with our findings.

In conclusion, ARI is a plastic developmental process controlled by crosslinked hormonal networks that trigger multilayered transcriptional cascades.

## Material and methods

### Plant material

Seeds of the *ninja-1myc2-322B* double mutant (Gasperini *et al*., 2015), *coi1-16* (Ellis and Turner, 2002), *35S:CKX1* (Werner *et al*., 2003), and *35S:RAP2-6L* (Krishnaswamy *et al*., 2011) lines were respectively provided by E.E. Farmer (University of Lausanne, Switzerland), L. Pauwels (VIB/PSB, Ghent, Belgium), T. Schmülling (Freie Universität Berlin, Germany) and N. Kav (University of Alberta, Edmonton, Canada). The triple mutant *ninja-1myc2-322B-35S:CKX1* was generated by crossing *ninja-1myc2-322B* and *35S:RAP2-6L* plants. The *Arabidopsis thaliana* ecotype Col-0 was used as the wild type and background for all the mutants and *35S:CKX1* transgenic line, except the *35S:RAP2.6L* transgenic line, which was in Ws-4 background.

### Adventitious and lateral root phenotyping and growth conditions

Seedlings were grown in adventitious root-inducing conditions as previously described (Sorin *et al*., 2005; Gutierrez *et al*., 2009, 2012; Lakehal *et al*., 2019*a*). Briefly, they were grown in dark conditions until their hypocotyls were 6-7 mm long then transferred to long day conditions. Numbers of AR (primordia and emerged) were counted under a binocular stereomicroscope 7 days after transferring the seedlings to light conditions. Numbers of emerged lateral roots were counted on the same day from scanned plates. The seedlings’ primary root lengths were measured from scanned plates using imageJ software (Schindelin *et al*., 2012), and their lateral root densities (LR numbers to primary root length ratios) were calculated.

### Gene expression experiments

#### Sample preparation

To check the effect of JA on *CKX1* expression, total RNA was extracted from whole wild type (Col-0) and mutant *coi1-16* seedlings, which were grown under long day conditions for five days post-germination then moved to sterile liquid media for overnight acclimation. The next day, the seedlings were treated with either jasmonic acid (J2500, Sigma) at selected concentrations or mock.

To characterize *CKX1* expression in the *ninja-1myc2-322B* double mutant and Col-0 seedlings during ARI, total RNA was extracted from dissected hypocotyls of etiolated seedlings. The seedlings were grown under AR-inducing conditions as described above, i.e. grown in the dark until their hypocotyls were 6-7 mm long (T0) then transferred to light for either 9 h (T9) or 24h (T24).

To examine effects of CK and JA on *RAP2.6L* expression, total RNA was extracted from etiolated wild type (Col-0) seedlings. As *RAP2.6L* expression is downregulated by light, we decided to check its expression in dark-grown seedlings. The seedlings were first etiolated in the dark until their hypocotyls were approximately 6 mm long then transferred to liquid media for acclimation overnight. The next day they were treated with *t*Z, JA, *t*Z+JA (at concentrations listed above) or mock.

#### RNA isolation and cDNA synthesis

Total RNA was extracted using a RNAqueous® Total RNA Isolation kit (Ambion™) then treated with DNaseI using a DNA*free* Kit (Ambion™) to remove contaminating DNA. cDNA was synthesized by reverse-transcription of the RNA using a SuperScript II Reverse transcriptase kit (Invitrogen) with anchored-oligo(dT)18 primers. All the steps were performed according to the kit manufacturers’ instructions.

#### Quantitative RT-PCR (qRT-PCR)

Transcript amounts were quantified by qRT-PCR, using triplicate reaction mixtures containing 5 μL of cDNA, 0.5 μM of both forward and reverse primers, and 1× LightCycler 480 SYBR Green I Master (Roche) (final volume, 20 μL) according to the manufacturer’s instructions. All these experiments were performed with at least two independent biological replicates. Relative amounts of transcripts of the target genes were calculated as previously described (Gutierrez *et al*., 2009) and regarded as significantly down- or up-regulated if fold differences were ≤ 0.75 or ≥ 1.5, respectively, with *P*-values ≤ 0.05. Reference genes were validated as the most stably expressed genes under our experimental conditions (Gutierrez *et al*., 2009) using GenNorm software and the most stable pair were used to normalize the quantitative PCR data. Data obtained using *TIP41* as the reference gene are presented here. Sequences of primers used for all target and reference genes are listed in supplementary Table 7.

#### CK quantification

For CK quantification, wild type seedlings were grown under AR-inducing conditions as described above. The seedlings were first etiolated in the dark until their hypocotyls were 6-7 mm long (T0) then transferred to light for either 9 h (T9) or 72 h (T72). Etiolated hypocotyls were collected and rapidly dried on tissue paper then frozen in liquid nitrogen. The samples were stored at -80°C until further use. Samples were prepared from six biological replicates, each consisting of two technical replicates. Cytokinin species (ribotides, ribosides, free-bases and glucosyl conjugates) were quantified as previously described (Lakehal *et al*., 2020*a*) from 20 mg fresh weight samples using published methodology (Svačinová *et al*., 2012; Antoniadi *et al*., 2015).

## Author contributions

AL, AD and CB conceived and designed the research; AL, AD and ON performed the research; AL, AD, ON analyzed the data, AL, AD and CB wrote the manuscript; CB and ON acquired the funding.

## Acknowledgements

The authors sincerely thank Hana Martínková and Petra Amakorová for their help with phytohormone analyses. This work was supported by grants from the Swedish Research Council (VR), the Swedish Research Council for Research and Innovation for Sustainable Growth (VINNOVA), the K&A Wallenberg Foundation, the Carl Trygger Foundation, and the Carl Kempe Foundation awarded to CB, together with grants from the Ministry of Education, Youth and Sports of the Czech Republic (European Regional Development Fund-Project ‘Plants as a tool for sustainable global development’ no. CZ.02.1.01/0.0/0.0/16_019/0000827), and the Czech Science Foundation (Project no. 19-00973S) awarded to ON

