## Supplemental Information for "Jasmonate inhibits adventitious root initiation through transcriptional repression of *CKX1* and activation of *RAP2.6L* transcription factor in Arabidopsis"

**The following Supporting Information is available for this article:**

**Supplementary figures**

**Figure S1**: Dark-light transition causes changes in DHZ in etiolated hypocotyls.
**Figure S2:** MeJA negatively regulates *CKX1* expression.
**Figure S3:** *CKX1* expression during ARI.

**Supplementary tables**

**Table S1-S3:** list of differentially expressed genes (DEGs) in the wild type (Col-0) during adventitious root initiation (excel document)
**Table S4-S6:** list of differentially expressed transcription factors (TFs) in the wild type (Col-0) during adventitious root initiation (excel document)
**Table S7:** Primers used for qRT-PCR in this study

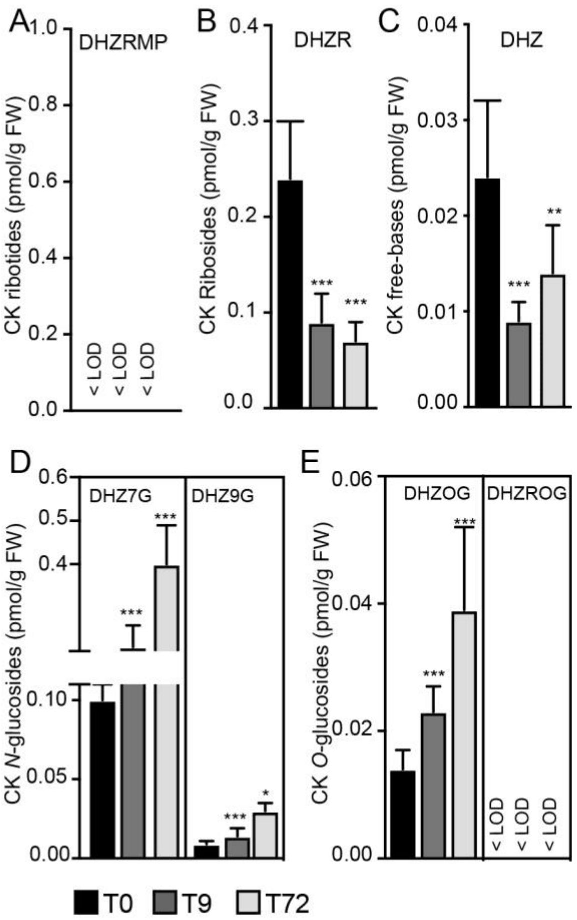


**Figure S1**: Dark-light transition causes changes in DHZ in etiolated hypocotyls.
CK nucleotides **(A),** CK ribosides **(B)** CK bases **(C),** CK N-glucosides **(D)** and CK O-glucosides **(E)** quantified in hypocotyls dissected from Col-0 seedlings grown in the dark until their hypocotyls were 6 mm long then shifted to the light for 9 h (T9) and 72 h (T72). Means and standard deviations**.** Asterisks indicate significant differences at T9 and T72 versus T0 according to Analysis of Variance with t-tests: ***,** **, and *** 0.05 > $p$ > 0.01, 0.01 > $p$ > 0.001, and $p$ < 0.001, respectively). <LOD means under the limit of detection.

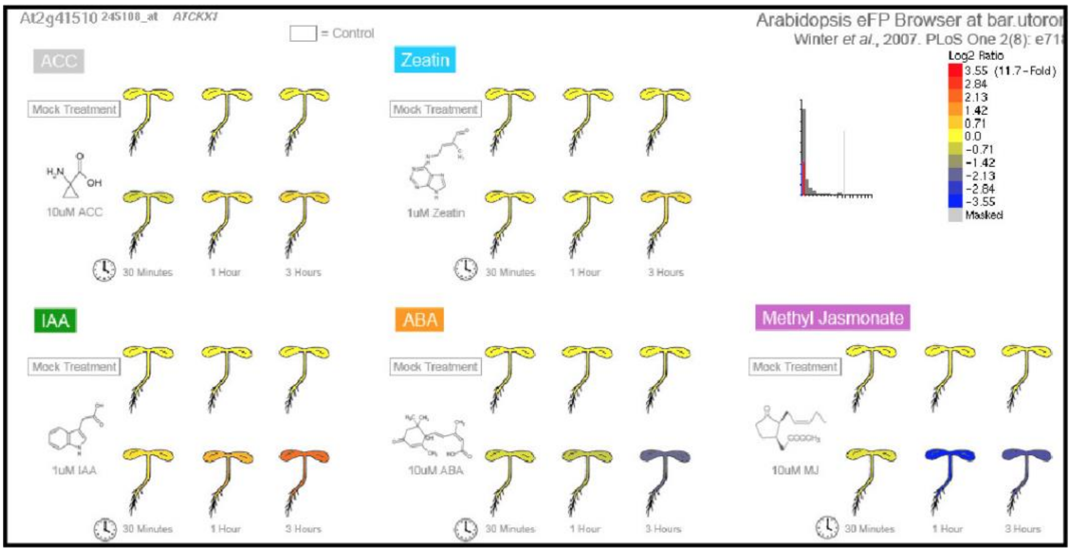


**Figure S2:** MeJA negatively regulates *CKX1* expression. Data were retrieved from http://bar.utoronto.ca.

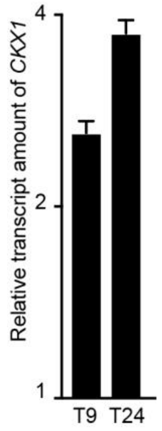


**Figure S3:** *CKX1* expression pattern during ARI.
Relative amounts of *CKX1* transcripts quantified by qRT-PCR. Amounts of transcripts extracted from dissected hypocotyls of wild-type seedlings etiolated in the dark until the hypocotyl was 6 mm long (T0) then moved to the light for 9 h (T9) or 24 h (T24) relative to amounts detected at T0. Means (bars) and standard errors of the mean (whiskers) obtained from three technical replicates. The experiment was repeated with another independent biological replicate and gave similar results.

**Table S7: Primers used for qRT-PCR in this study**

| Primer name | Gene number | Primer forward | Primer reverse |
|---|---|---|---|
| qRT-RAP2.6L | AT5G13330 | CAAGGCCCTACTACCACCACAA | GGTCGAGGAGGAGGTGAGTTC |
| qRT-CKX1 | AT2G41510 | ATGGATCAGGAAACTGGCAA | AGATGAAAACAAAGTGGATGGAA |
| qRT-TIP41 | At4g34270 | GCTCATCGGTACGCTCTTTT | TCCATCAGTCAGAGGCTTCC |
